# New non-bilaterian transcriptomes provide novel insights into the evolution of coral *skeletomes*

**DOI:** 10.1101/594242

**Authors:** Nicola Conci, Gert Wörheide, Sergio Vargas

**Affiliations:** Department of Earth and Environmental Sciences, Palaeontology & Geobiology, Ludwig-Maximilians-Universität, Munich, Germany; GeoBio-Center LMU, Ludwig-Maximilians-Universität, Munich, Germany; SNSB - Bayerische Staatssammlung für Paläontologie und Geologie, Munich, Germany

**Keywords:** coral calcification, biomineralization, Octocorallia, galaxin, molecular evolution, Scleractinia

## Abstract

A general feature of animal *skeletomes* is the co-presence of taxonomically widespread and lineage-specific proteins that actively regulate the biomineralization process. Among cnidarians, the skeletomes of scleractinian corals have been shown to follow this trend, however in this group distribution and phylogenetic analyses of biomineralization-related genes have been often based on limited numbers of species, with other anthozoan calcifiers such as octocorals, being overlooked. We *de-novo* sequenced the transcriptomes of four soft-coral species characterized by different calcification strategies (aragonite skeleton *vs*. calcitic sclerites) and data-mined published non-bilaterian transcriptomic resources to construct a taxonomically comprehensive sequence database to map the distribution of scleractinian and octocoral *skeletome* components. At the large scale, no protein showed a ‘Cnidaria+Placozoa’ or ‘Cnidaria+Ctenophora’ distribution, while some were found in cnidarians and Porifera. Within Scleractinia and Octocorallia, we expanded the distribution for several taxonomically restricted genes (TRGs) and propose an alternative evolutionary scenario, involving an early single biomineralization-recruitment event, for galaxin *sensu stricto*. Additionally, we show that the enrichment of acidic residues within skeletogenic proteins did not occur at the Corallimorpharia-Scleractinia transition, but appears to be associated with protein secretion in the organic matrix. Finally, the distribution of octocoral calcification-related proteins appears independent of skeleton mineralogy (*i*.*e*. aragonite/calcite) with no differences on the proportion of shared skeletogenic proteins between scleractinians and aragonitic or calcitic octocorals. This points to *skeletome* homogeneity within but not between groups of calcifying cnidarians, although some proteins like galaxins and SCRiP-3a could represent instances of commonality.

## 1. Introduction

Cnidaria is a phylum of marine and freshwater invertebrates currently comprising ∼9,000 valid species. Their synapomorphy is the cnidocyte, a unique cell type used for locomotion and prey capture (Holstein 1981; Kass-Simon et al. 2002). Cnidarians have been important reef-building organisms throughout Earth history (Wood 1999) and are the main ecosystem engineers in today’s coral reefs (Wild et al. 2011). Several taxa produce a rigid mineral skeleton made of calcium carbonate (CaCO_3_) and those are found in the anthozoan order Scleractinia and the subclass Octocorallia, as well as in the hydrozoan families of Milleporidae, Stylasteridae and Hydractiniidae. Calcification apparently has evolved multiple times independently within Cnidaria (*i*.*e*. in scleractinians; Romano and Cairns 2000) and hydractinians (Miglietta et al. 2010), and according to molecular clock estimates the origin of the capacity to calcify arose prior to the appearance of cnidarian skeletons in the fossil record (Cartwright and Collins 2007; Erwin et al. 2011; Van Iten et al. 2016).

A common feature of most calcifying organisms is their ability to biologically control and regulate the formation of their skeletons. Although the degree of such control in cnidarians is still debated and the underlying molecular mechanisms are not entirely understood (see Tambutté et al. 2011 for a review), two main regulatory mechanisms have been described. The first concerns the transport, availability and concentration of required ions, and involves proteins like carbonic anhydrases (Jackson et al. 2007; Moya et al. 2008; Bertucci et al. 2011) and bicarbonate transporters (Zoccola et al. 2015). The second putatively involves the skeletal organic matrix (SOM), an array of proteins (Puverel et al. 2004), acidic polysaccharides (Goldberg 2001) and lipids (Farre et al. 2010) occluded within the mineral fraction of the skeleton (Farre et al. 2010). Skeletal organic matrix proteins (SOMPs) have been suggested to play a role in the promotion or the inhibition of crystal growth (Allemand et al. 1998; Clode and Marshall 2003; Puverel et al. 2005) and in the regulation of mineral polymorphism (Goffredo et al. 2011) and are collectively referred to as the ‘skeletogenic proteins’ (Jackson et al. 2007), ‘biomineralization toolkits’ (Drake et al. 2013) or ‘skeletomes’ (Ramos-Silva et al. 2013) (Goffredo et al. 2011). The characterization of SOMPs and the study of their evolutionary history is thus essential to unravel the appearance and evolution of biomineralization also in cnidarians.

The first protein described and characterized from a coral skeleton was isolated from the organic matrix of the scleractinian coral *Galaxea fascicularis* and thus named galaxin (Fukuda et al. 2003). Galaxins are ubiquitous among scleractinians and putative homologs have been identified in several animal groups, including polychaetes (Sanchez et al. 2007), molluscs (Heath-Heckman et al. 2014) and sea urchins (Sodergren et al. 2006). Although structural similarities with vertebrate usherin (Bhattacharya et al. 2004) led to the proposition of an interaction between galaxin and *type IV* collagen (Bhattacharya et al. 2016), the role of galaxin in cnidarian skeletogenesis remains to be fully resolved (Bhattacharya et al. 2016). Following the first descriptions of single skeletogenic proteins, the advent of tandem mass-spectrometry allowed the simultaneous characterization of several proteins, offering a general overview of coral skeletal proteomes. To date, the proteome of three scleractinian corals: the two acroporids *Acropora digitifera* (Takeuchi et al. 2016) and *Acropora millepora* (Ramos-Silva et al. 2013), and the pocilloporid *Stylophora pistillata* (Drake et al. 2013) have been characterized.

The most abundant fraction of the coral skeletomes so far characterized is represented by acidic proteins (Ramos-Silva et al. 2013; Takeuchi et al. 2016), which supposedly drive crystal nucleation and growth (Wheeler et al. 1981; Addadi et al. 1987). Six acidic proteins have been described from the skeleton of *A*. *millepora* and two from *S*. *pistillata*. These include skeletal aspartic acid rich proteins (SAARPs) (Ramos-Silva et al. 2013) and secreted acidic proteins (SAPs) (Shinzato et al. 2011) - both found in *Acropora* - and two *S*. *pistillata* coral acid rich proteins (CARP4 and CARP5). Interestingly non-acidic regions of SAARPs match sequences found in other non-biomineralizing cnidarians and bivalves, making the high occurrence of acidic residues a potential secondary modification linked to biomineralization (Takeuchi et al. 2016).

Surveys of cnidarian transcriptomes and genomes have in fact revealed that only a small proportion of SOMPs in *A*. *millepora* appears to be taxonomically restricted genes (TRGs) in corals (Ramos-Silva et al. 2013), while the majority of SOMPs (ca. 80% in *A*. *millepora*) have putative homologs in non-calcifying cnidarians, like sea anemones and/or *Hydra magnipapillata* (Ramos-Silva et al. 2013). Also a recent transcriptome survey of corallimorpharians, skeleton-lacking cnidarians closely related to Scleractinia, has further shown that only six skeletogenic proteins appear to be taxonomically restricted to scleractinian corals (Lin et al. 2017).

So far, however, genomic and transcriptomic surveys have mainly focused on comparisons between scleractinian corals and a limited set of non-calcifying cnidarians (*e*.*g*. sea anemones, corallimorpharians and *Hydra*), systematically overlooking octocorals and calcifying hydrozoans (but see Guzman et al. 2018). Thus, very little information is currently available on the distribution of SOMPs across and within different lineages of calcifying cnidarians and consequently the evolutionary history of their biomineralization-related genes remains largely unexplored.

Here, we conducted an analysis of the distribution of coral *biomineralization toolkit* components across Anthozoa. This allowed us to trace the appearance of skeletogenic protein homologs and investigate observed differences between (*e*.*g*. Scleractinia *vs*. Octocorallia) and within lineages. In addition, we also compared biomineralization gene repertoires between and within 1) calcifying cnidarians and sponges displaying different calcification strategies (*i*.*e*. aragonite *vs*. calcite deposition, exoskeleton *vs*. endo-sclerites) like octocorals and scleractinians or calcareous sponges and the aragonitic demosponge *Vaceletia* sp. and 2) between them and their non-calcifying close relatives. For this we *de novo sequenced* the transcriptomes of four octocoral species, namely the massive, aragonitic blue coral *Heliopora coerulea*, and calcite producing species *Pinnigorgia flava, Sinularia* cf. *cruciata* and *Tubipora musica*, three sclerites-forming octocorals. These species cover all calcification strategies within Octocorallia. Data-mining of newly generated and publicly available sequence resources was then used to produce fine-scaled phylogenies for selected targets of interest including acidic proteins (e.g. CARPs, SAARPs), galaxin and carbonic anhydrases. These results contribute to our understanding of the functional diversity and evolutionary history of coral skeletomes.

## 2. Material and Methods

### Generation of octocorals reference transcriptomes

To obtain reference transcriptomes for our target octocoral species, samples of *H*. *coerulea, T*. *musica, P*. *flava* and *S*. cf. *cruciata*, were mechanically collected from colonies cultured in the aquarium facilities of the Chair for Geobiology & Paleontology of the Department of Earth-and Environmental Sciences at Ludwig-Maximilians-Universität München in Munich (Germany) and kept under control conditions (temperature 25.1 ± 0.5°C, pH 8.2 ± 0.1) for ca. 1 month before fixation in liquid nitrogen and subsequent storage at −80°C.

For RNA extraction the samples were homogenized in 1-2 ml TriZol (Thermofisher) using a Polytron PT Homogenizer (Kinematica), and subsequently centrifuged (20 min at 13,500 rpm and 4°C) to remove remaining skeletal debris. A modified TriZol protocol (Chomczynski and Mackey 1995) was used for RNA purification and the concentration and integrity of the extracted RNA were assessed on a NanoDrop 2100 spectrophotometer and a Bioanalyzer 2100 (Agilent), respectively. For each species, RNA samples with a RIN > 8.5 were used to prepare strand-specific libraries that were paired-end sequenced (50 bp reads) on an Illumina HiSeq 2000 sequencer at the EMBL Core Center in Heidelberg (Germany). For *H*. *coerulea*, additional strand-specific libraries were generated with the SENSE mRNA-Seq Library Prep Kit V2 for Illumina (Lexogen), and sequences on an Illumina NextSeq 500 at the Kinderklinik und Kinderpoliklinik im Dr. von Haunerschen Kinderspital.

Quality control of assembled reads was done with FastQC (www.bioinformatics.babraham.ac.uk) and low quality reads (Q<28) were removed with the Filter Illumina program from the Agalma-Biolite transcriptome package (Dunn et al. 2013). In addition, reads were mapped against a set of microbial genomes with Bowtie 2 (Langmead and Salzberg 2012) and mapping reads were discarded. Transcriptome assembly was performed with Trinity v.2.5.1 (Grabherr et al. 2011). Contigs with a length <300 bp were discarded. Transcriptome completeness was assessed with BUSCO 3.0.2 (Simão et al. 2015) using the Metazoa odb9 dataset and protein sequences were predicted with TransDecoder v.3.0.1. Summary statistics for each assembly is provided in Table 1. The bioinformatic workflow used is available at https://galaxy.palmuc.org. Reads were deposited at the European Nucleotide Archive (https://www.ebi.ac.uk/ena) under Accession Number PRJEB30452. Assemblies, untrimmed/trimmed alignments, and output tree files from the various analyses are available at https://gitlab.lrz.de/palmuc/concietal_octoskeletomes

**Tab 1.**
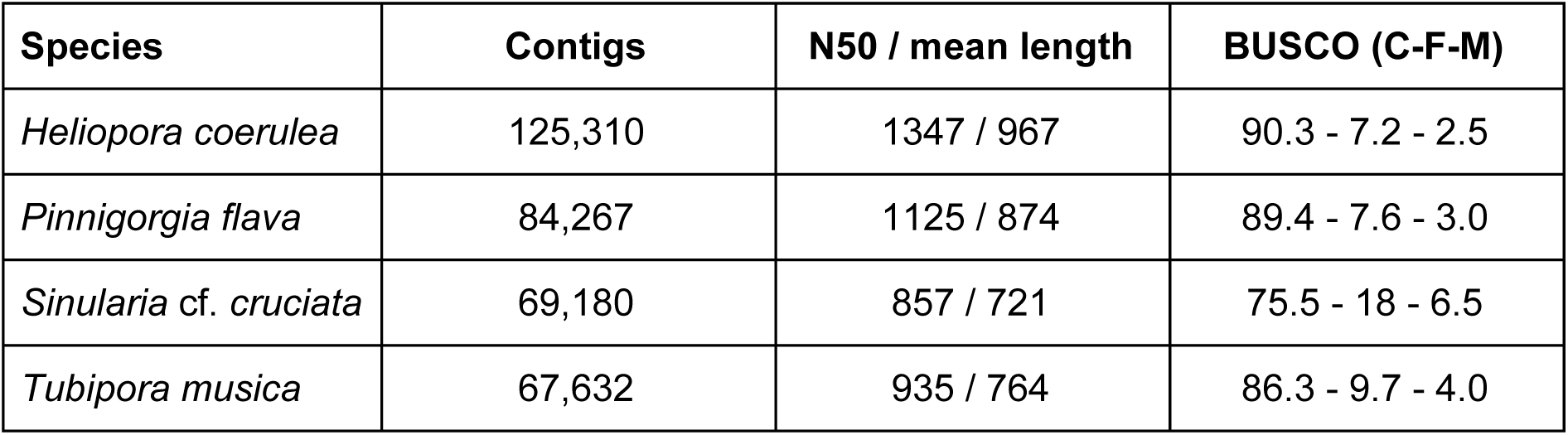
Summary statistics for the assembled meta-transcriptomes. For BUSCO analysis, percentages of complete (C), fragmented (F) and missing (M) orthologs are provided,

### Database construction and homologs search/analysis

To construct the homolog database (S.Mat1) of calcification-related proteins, newly assembled transcriptomes were added to a sequences database of representative of the non-bilaterian metazoan phyla Cnidaria, Porifera, Placozoa, and Ctenophora. To construct the database, publicly available resources for target organisms (excluding tissue-specific transcriptomes) were uploaded on our local Galaxy server (https://galaxy.palmuc.org). Source details for each dataset is provided in S.Mat1. When protein sequences were available, these were directly converted to a protein Blast database (makeblastdb). Nucleotide sequences were first translated with TransDecoder Galaxy Version 3.0.1 (Haas et al. 2013). For cnidarians, Blast databases were individually searched (BlastP, e-value cut-off > 1e^−09^) to retrieve putative homologs of coral calcification-related sequences. For Porifera, Ctenophora and Placozoa, databases provided in Eitel et al. (2018) were searched using the same criteria listed above. Search queries (S.Mat1) included scleractinian skeletogenic proteins from *A*. *millepora* (Ramos-Silva et al. 2013) and *S*. *pistillata* (Drake et al. 2013), and small cysteine rich proteins (SCRiPs) from *Orbicella faveolata* (Sunagawa et al. 2009). From *S*. *pistillata*, two additional SAARP-like acidic proteins that were included in the phylogenetic analysis in Bhattacharya et al. (2016) were additionally used as search queries. Octocoral queries comprised carbonic anhydrases from both *Corallium rubrum* (Le Goff et al. 2016) and *Lobophytum crassum* (Rahman et al. 2006) and scleritin (Debreuil et al. 2012). Features including sequence length and amino acid composition of identified homologs were determined with ProtParam (Gasteiger et al. 2005). To predict the presence of signal peptide, transmembrane regions, and protein domains, SignalP 4.1 (Petersen et al. 2011), TMHMM 2.0 (Sonnhammer et al. 1998) and InteProScan (Jones et al. 2014) were used, respectively.

### Homolog selection for phylogenetic analysis

For the phylogenetic reconstruction of acidic proteins, all best-hit sequences identified through the Blast search described above were used. Additionally, non-scleractinian sequences retrieved after Blast analyses were used as query against scleractinian datasets (using BlastP, e-value < 1e^−09^) (S.Mat1). If the corresponding scleractinian best-hit differed from those identified using the previous query, the sequences were also considered for phylogenetic analysis. The analyses of galaxin *sensu stricto* (scleractinian orthologs of *G*. *fascicularis* galaxin) and galaxin-related proteins (*i*.*e*. other putative homologs within and outside scleractinians) are based on all putative homologs (e-value < 1e^−09^), with the exception of those matching galaxin-like 1 and 2 (ADI50284.1 and ADI50285.1), as these are exclusively expressed during early stages of calcification (Reyes-Bermudez et al. 2009). Predicted, partial sequences of less than 200 aa long were excluded. In addition to scleractinians, we surveyed taxa in which galaxin related proteins have been identified, namely from Mollusca, Annelida (Class Polychaeta), and Echinodermata. All resulting sequences were searched, using BlastP, (e-value < 1e^−09^) against the NCBI non-redundant database to avoid including usherin homologs in the dataset. Homologous sponge collagen IV sequences were searched using the *type IV* collagen (Q7JMZ8) identified in the homoscleromorph sponge *Pseudocoriticium jarrei* as query. Analysis was limited to the N-terminal NC1 domain. Sequences of each putative homolog were checked for the presence of conserved cysteines (Aouacheria et al. 2006) and added to the collagen IV-spongins dataset in Aouacheria et al. (2006). Finally octocoral homologs for the carbonic anhydrases (CA) CruCA1-6 (Le Goff et al. 2016) were searched in all octocoral datasets considered and added to the CAs dataset used in Voigt et al. (2014)

### Phylogenetic analysis

Protein sequences were aligned with MAFFT (Katoh and Standley 2013) and MUSCLE (Edgar 2004) to investigate possible effects of the aligning algorithm on the final phylogeny. Alignment was followed by a first site selection with Gblocks (Castresana 2002) and final manual curation. Best-fit models were determined with Prottest 3 (Darriba et al. 2011). Maximum-likelihood and bayesian analyses were performed in PhyML 3.1 (Guindon et al. 2010) from Seaview 4 (Gouy et al. 2010) with 500 bootstrap replicates and MrBayes 3.2 (nruns=2, samplefreq=100; Huelsenbeck and Ronquist 2001; Ronquist et al. 2012), respectively. Effective Sample Sizes (EES >200) and burn-in fractions (0.20-0.25) were determined with Tracer v.1.6 (http://tree.bio.ed.ac.uk/software/tracer/).

## 3. Results

### Distribution analysis of skeletogenic proteins

The presence for coral skeletogenic protein homologs differed markedly between protein categories (Fig. 1). Carbonic anhydrases, peptidases and extracellular/adhesion proteins display the widest taxonomic distribution, although similarity was often limited to conserved domains within protein sequences. In sponges and cnidarians however, differences could also be observed in terms of domain presence. These include the zona pellucida, observed only in calcareous sponges, the absence of the MAM domain in Demospongiae, as reported in Riesgo et al. (2014), and the cupredoxin domain in Hydrozoa. In contrast, all secreted acidic proteins (SAPs) and all small cysteine rich (SCRiPs) proteins with the sole exception of SCRiP-3a (ACO24832.1), which was detected in Scleractinia and Octocorallia, showed the most taxonomically restricted distribution. Despite the presence of proteins found only among certain scleractinian families (*e*.*g*. SAPs, Threonine-rich protein), no protein hitherto isolated from the skeleton of *A*. *millepora* was found here restricted to acroporids. No protein exhibited a ‘Cnidaria+Placozoa’ or ‘Cnidaria+Ctenophora’ distribution pattern, while a small set of coral SOMPs appeared to possess homologs in Cnidaria and Porifera. These include galaxin-related proteins and the uncharacterized *A*. *millepora* protein USOMP-5 (B8VIU6.1). Although absent in Homoscleromorpha and Hexactinellida, galaxin-related proteins are ubiquitous among calcareous sponges and also found in all three currently described subclasses of Demospongiae. Within Heteroscleromoprha however, differences were observed between groups as no galaxin-related protein was retrieved from the genome of *Amphimedon queenslandica* (Srivastava et al. 2010), while a significant hit was returned from the genome of *Tethya wilhelma* (Francis et al. 2017). The highest occurrence rate for USOMP-5 homologs in sponges was observed in Homoscleromorpha, but matches were detected in all groups. Although no protein domain was originally reported for B8VIU6.1 in *A*. *millepora* (Ramos-Silva et al. 2013), analysis of matching sequences from sponges revealed the presence of fibrinogen-related subdomains (IPR014716, IPR036056) within the protein (S.Fig.1). Domain location partly overlaps the conserved region of the protein, and might thus explain the detected local similarity.

**Fig.1.**
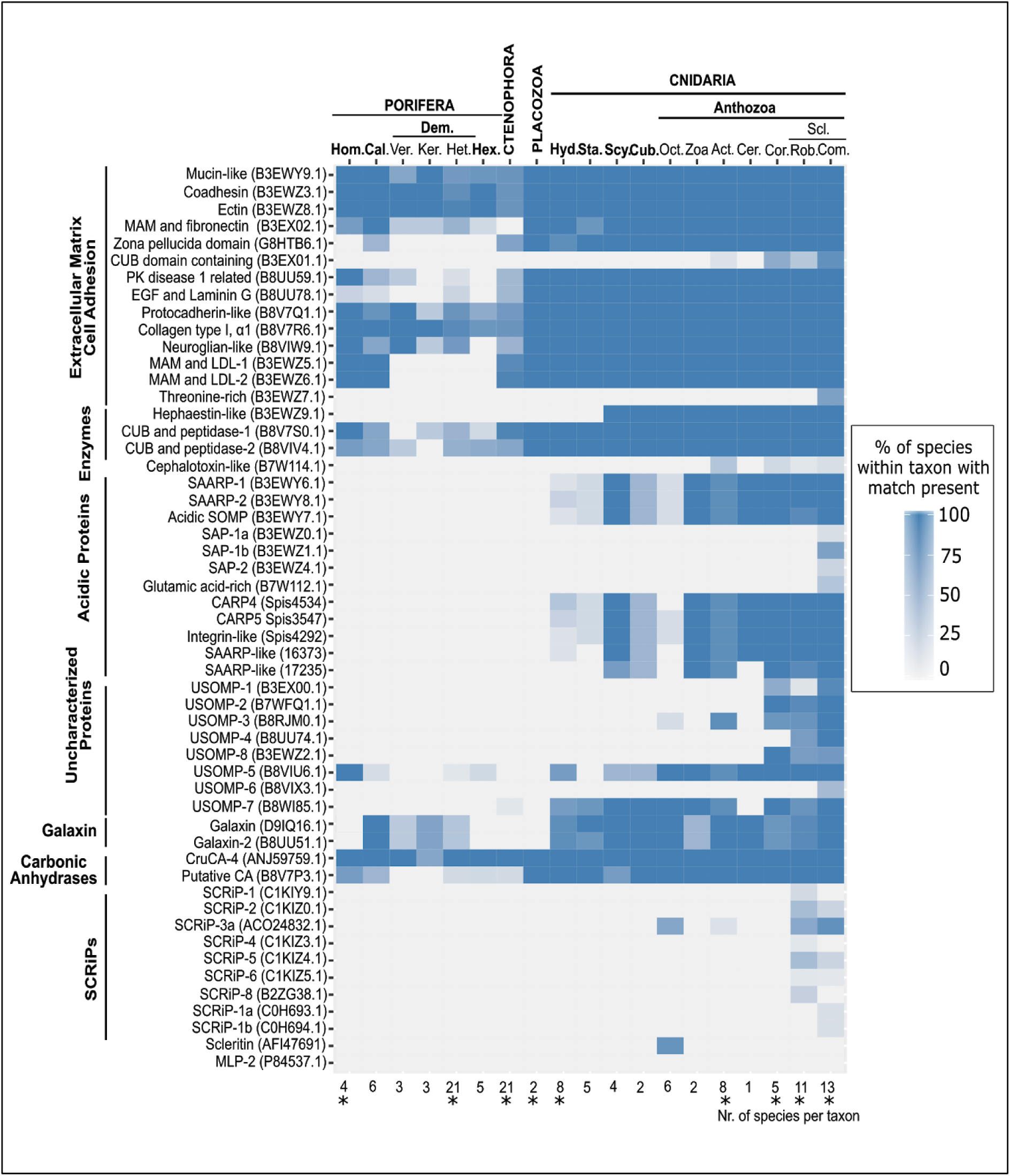
Pattern of presence of homologs (BlastP, e-value < 1e^−09^) of coral biomineralization-related protein across non-bilaterian metazoans. Lower x axis indicates number of species surveyed within a particular group. Asterisk: genomic data available for at least one species within the group. Protein categories ‘Extracellular matrix - Cell Adhesion’, ‘Enzymes’, ‘Uncharacterized Proteins’ and ‘Galaxins’ based on Ramos-Silva et al. (2013). Taxa in capital and bold: phyla. Taxa in bold: classes. Normal text: subclasses or lower taxonomic levels. Hom: Homoscleromorpha, Cal: Calcarea, Ver: Verongimorpha, Ker: Keratosa, Het: Heteroscleromorpha, Dem: Demospongiae, Hex: Hexactinellida, Hyd: Hydrozoa, Sta: Staurozoa, Scy: Scyphozoa, Cub: Cubozoa, Oct: Octocorallia, Zoa: Zoantharia, Act: Actinaria, Cer: Ceriantharia, Cor: Corallimorpharia, Scl: Scleractinia, Rob: Robusta (Scleractinia), Com: Complexa (Scleractinia).

Despite exhibiting different patterns, other skeletogenic protein homologs were restricted to Cnidaria. Acidic proteins SAARPs and CARPs produced significant Blast matches among several cnidarian groups, although the presence of acidic regions (*i*.*e*. sequences segments enriched in aspartic and glutamic acid) appears characteristic of scleractinian corals (see below). Members of the SAP acidic family were, on the other hand, detected solely in complex scleractinians, but not only in acroporids as previously suggested (Shinzato et al. 2011; Takeuchi et al. 2016). Homologs of SAP-1b (B3EWZ1.1) are in fact present within families Dendrophylliidae and Agariciidae.

Other uncharacterized proteins (USOMPs) displayed varying presence/absence patterns. USOMP-7 (B8WI85.1) and USOMP-3 (B8RJM0.1) were found across cnidaria and Anthozoa respectively. The latter also represents the only difference we detected between aragonitic and calcitic octocorals as this protein was solely found in *H*. *coerulea*. As reported in Lin et al. (2017), USOMP-1 is present in anemones and scleractinians, while both USOMP-2 and USOMP-8 first appear in corallimorphs. Finally, USOMP-4 and USOMP-6 are restricted to scleractinians, although the first is shared by complex and robust corals and the second was only found in the families Acroporidae and Agariciidae.

No significant match was detected among octocorals for the acidic carbonic anhydrase MLP-2 (Rahman et al. 2006), while we retrieved homologs across the group for both scleritin and five (CruCA1-5) of the six carbonic anhydrases described for *C*. *rubrum* (S.Fig8, S.Fig9), including the putative skeletogenic CruCA-4. No difference has thus been observed for octocoral calcification-related proteins between aragonite and calcite-deposing species.

### Phylogenetic analysis of acidic proteins

Phylogenetic analysis split acidic proteins and their non-acidic homologs into five well-supported clades: two of these (marked as “S” for skeletogenic clades) are occupied by proteins found occluded in coral skeletons. Only scleractinians are represented within these groups. S1 contains homologs for the acidic SOMP (B3EWY7) and P27 isolated from *A*. *millepora* (Ramos-Silva et al. 2013) and *S*. *pistillata* (Drake et al. 2013), respectively. Both these proteins display shorter acidic regions and a lower aspartic acid content compared to SAARPs and CARPs, which occupy clade S2. Tree topology within this group did however change between phylogenies obtained using different alignment methods (*i*.*e*. MUSCLE *vs*. MAFFT). In the MAFFT-based tree displayed below, (Fig.2a) CARPs and SAARPs split into two distinct subgroups although bootstrap support was low. All other sequences were divided in three non-skeletogenic (NS) clades. Taxonomic diversity for these groups did differ and ranged from Cnidaria (NS1) to scleractinians (NS3), while NS2 contained scleractinians and corallimorphs.

**Fig.2.**
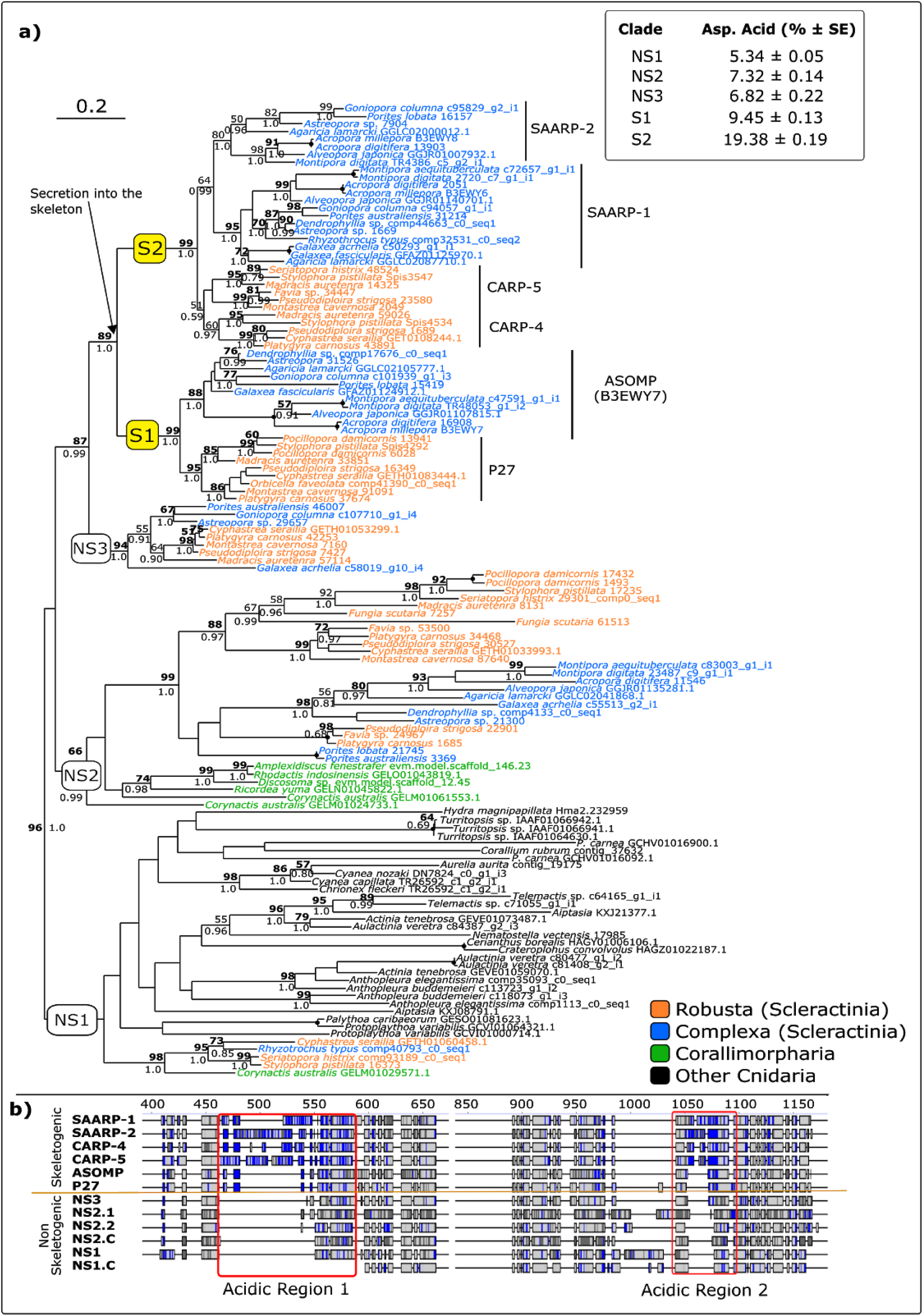
**a)** Phylogenetic tree (ML, 500 bootstrap replicates) of scleractinian acidic proteins and putative homologs in other cnidarian groups. Best-fit model: WAG+F+G+I. Tree displayed in figure based on protein sequences aligned with MAFFT. MUSCLE alignment and tree available in S.Fig4. Bold number: node supported (>50) also in MUSCLE phylogeny. Dot on node indicates full support (100% bootstrap, 1.0 posterior probability) in both phylogenies. Support for nodes with bootstrap < 50 not shown regardless of posterior probability value. Skeletogenic clades (S) (highlighted in yellow) include acidic proteins found in coral skeletons (Drake et al. 2013; Ramos-Silva et al. 2013). NS (non-skeletogenic) clades: acidic proteins not extracted from coral skeletons. **b)** Consensus sequences (60%) alignment for each clade. Alignment shows the position and distribution of acidic residues (aspartic and glutamic acid) highlighted in blue. Light gray: other conserved residues. Dark gray: non-conserved residues. Complete alignment available S.Mat2. When corallimorph sequences were present in a clade, these were analysed separately to highlight difference with scleractinian proteins Corallimorph consensus sequences IDs end in “.C”. NS2 clade was split into NS2.1 (includes *P*. *australiensis* 3369, *P*. *lobata* 21745, *Favia* sp. 24967, *P*. *carnosus* 1685 and *P*. *strigosa* 22901) and NS2.2 (all other scleractinian sequences) because the position of NS2.1 was not congruent between phylogenies and was also retrieved as sister group to the rest of NS2 scleractinian proteins (S.Fig3). Top right corner: mean (± SE) content (%) of aspartic acid within acidic proteins. Average estimated on predicted complete sequences only.

When aligned with MUSCLE, SAARP-2 grouped with both CARPs but support was again low (S.Fig3). The internal topology of clade NS2 was also affected. When aligned with MUSCLE both *Porites* sequences, together with *Flavia* sp. 24967, *Platygyra carnosus* 1685 and *Pseudodiploria strigosa* 22901, were placed as sister group to other scleractinians (S.Fig.3). The split between corallimorphs and scleractinians within NS2 was nevertheless retrieved in both phylogenies. All other cnidarian sequences grouped with the scleractinian homologs of *S*. *pistillata* protein 17235 (NS1). As previously reported (Takeuchi et al. 2016), similarity between acidic proteins and their putative homologs is restricted to non-acidic regions. Although the transition between non-skeletogenic to skeletogenic proteins is not marked by severe increases in terms of aspartic acid content (6.82% in NS3 *vs*. 9.45% in S1), analyses of the consensus sequences for the different clades obtained here showed that aspartic and glutamic acid accumulation takes place only within acidic regions. The increase in aspartic/glutamic acid content and the appearance of acidic regions correspond with the secretion of the protein into the skeleton matrix and not with the shift between corallimorphs and scleractinian sequences (Fig.2b). Within B3EWY7-P27 the increment appears restricted to the first acidic region, and it then continues in SAARP1 and CARP4, ultimately escalating in SAARP-2 and CARP-5 which exhibit the longest extension of the first acidic region.

### Galaxin and *type IV* Collagen

Phylogenetic analysis of metazoan galaxin-related proteins revealed high degrees of polyphyly among lineages both at the phylum and lower levels, with only terminal nodes displaying moderate to high support (Fig.3). Taxonomically uniform clades can be observed in either the MAFFT or MUSCLE-based phylogeny. These included galaxin-related proteins from calcareous sponges, octocorals and Hydrozoa. However, for the vast majority of these clades, both support and topology appeared susceptible to the alignment algorithm employed.

**Fig.3.**
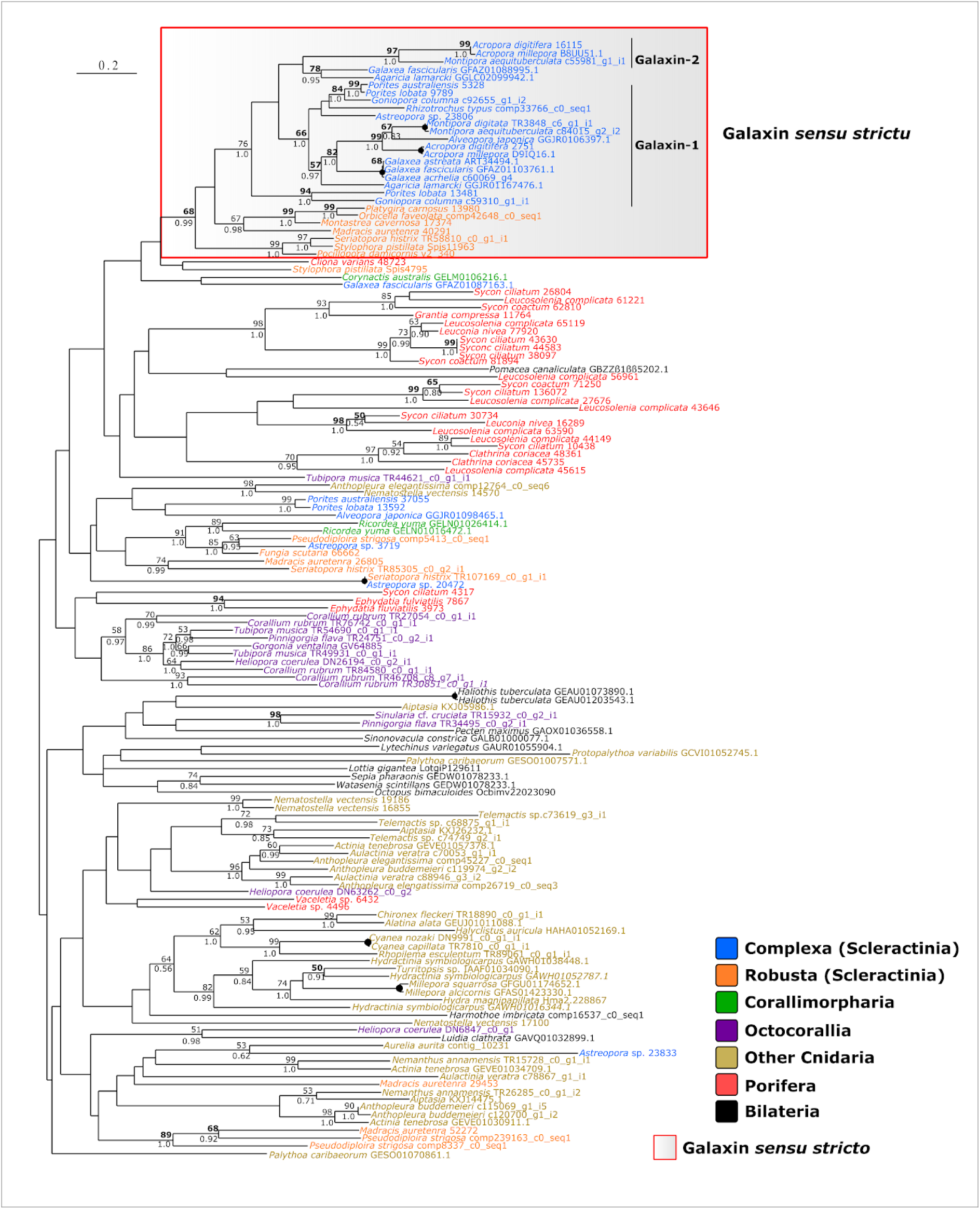
Phylogenetic analysis (ML; 500 bootstrap replicates) of metazoan galaxin-related proteins. Tree displayed in figure based on protein sequences aligned with MAFFT. MUSCLE alignment and tree available in S.Mat6. Bold number: node supported (>50) also in MUSCLE phylogeny. Dot on node indicates full support (100 bootstrap, 1.0 posterior probability) in both phylogenies. Support for nodes with bootstrap < 50 not shown regardless of posterior probability value.

Exception to this general pattern is a scleractinian-only clade comprising both complex and robust corals. The group includes both *A*. *millepora* skeletogenic (D9IQ16.1 and B8UU51.1) and the original *G*. *fascicularis* galaxins. The unifying feature of this clade is the RXRR endoprotease target motif described in Fukuda et al. (2003) (S.Fig.4). This RXRR motif is not unique to scleractinians, but it was not detected in any other galaxin-related protein within the group. Its presence thus appears to effectively discriminate a group of galaxins, here dubbed galaxins *sensu stricto*, from galaxin-related proteins.

Although the monophyly of galaxins *sensu stricto* was robust to the alignment algorithm, its internal topology was affected, with galaxin-2s and *R*. *typus* sequences nesting either within Complexa (MAFFT) or Robusta (MUSCLE). When performing the analysis on galaxin *sensu stricto* sequences only, galaxin-2 sequences concordantly grouped together with other complex scleractinians (S.Fig 3), in agreement with the topology derived from the MAFFT alignment and presented in Fig. 3. To investigate putative interactions between galaxin related proteins and collagen IV, we mapped the distribution of both proteins in Porifera, as both are present but not ubiquitous in the phylum (Fig.4).

**Fig.4.**
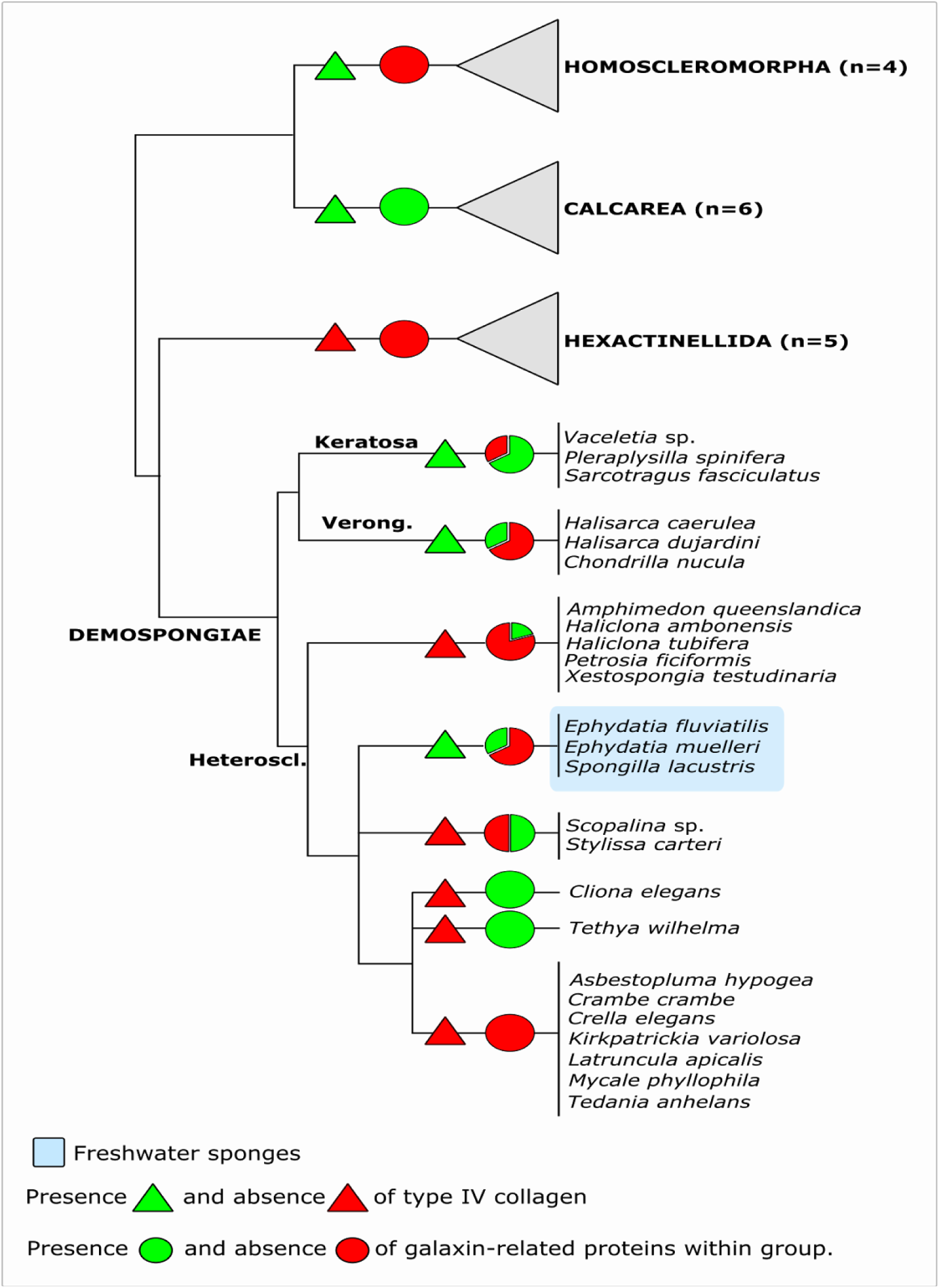
Presence-absence analysis of *type IV* collagen and galaxin-related proteins within Porifera. For galaxin-related proteins, data is presented as percentage of species within group in which one significant match (BlastP, e-value < 1e^−10^) was detected. When present, collagen IV was found in all species considered for a particular taxon (S.Mat 1). Phylogenetic relationships between sponge classes based on Simion et al. (2017). Phylogeny of Demospongiae based on Morrow et al. (2015). Heteroscl: Heteroscleromorpha. Verong: Verongimorpha.

As for galaxin-related proteins, *type IV* collagen is present across calcareous sponges, while Homoscleromorpha are the only sponge class with collagen IV but no galaxin homologs. The protein is present in both keratose and verongimorph sponges, while within Heteroscleromorpha it appears associated with the freshwater environment. Finally, neither protein is present in glass sponges (Hexactinellida).

Both phylogenetic analysis resulted in monophyly of collagen IV for all three sponge classes in which the protein is present (S.Fig 6-7). In one instance (MAFFT based phylogeny) support for monophyly of Porifera was also retrieved.

## 4. Discussion

A common feature of skeletal proteomes is the presence of both taxonomically widespread components possessing homologs in other, not necessarily calcifying, organisms and of lineage-specific innovations or taxonomically restricted genes (TRGs) (Ramos-Silva et al. 2013; Kocot et al. 2016). The diversity of evolutionary histories, characterising skeletogenic proteins, hence render the need of phylogenetic analyses and gene distribution maps essential steps to examine the evolution of biomineralization. A key requisite to conduct such assessments is in turn the need for extensive taxon sampling. Here we datamined available resources across non-bilaterian metazoans to examine the distribution of skeletogenic proteins, allowing comparative investigations of the genetic repertoires of diverse calcifying organisms, and produced detailed phylogenies for key components of coral *biomineralization toolkits*.

Distribution analysis reflected evolutionary heterogeneity, with homologs presence ranging from across phyla to selected families. Although a few coral skeletogenic proteins remain largely restricted taxonomically, increased taxon sampling resulted in the expansion of the taxonomic distribution for several *toolkit* components. In these cases, the most common pattern was the presence across phyla, or limited but conserved within Cnidaria or Scleractinia, which points against the involvement of these proteins in biomineralization across groups. SCRiP-3a and galaxin-related proteins are, however, potential targets for future (functional) research, because of their presence pattern (*e*.*g*. SCRiP-3a found among calcifying anthozoans only). The distribution map of the latter within sponges is of particular interest as it has been shown that these proteins are present in all calcifying species, regardless of their taxonomic position. Moreover, the presence of multiple potential galaxin homologs among Calcarea and their absence among homoscleromorphs and glass sponges, supports an involvement in calcium carbonate biomineralization, while the distribution of galaxin-related proteins in Demospongiae is less clear. Although the apparent conflict in genomic information could reflect genuine differences between sponge clades, the current absence of defining features within galaxin-related proteins does not allow to rule out wrong homology inferences that may add noise to our alignments and negatively influence phylogenetic inference.

As for galaxin-related proteins, collagen IV appears either ubiquitous or absent in different sponge classes, while a patchy distribution can be observed among groups of Demospongiae. Within Heteroscleromorpha presence of *type IV* collagen appears however, as previously hypothesized by Riesgo et al. (2014), associated with the freshwater environment, but among keratose sponges it could be related to the collagenous framework of their organic skeletons (Junqua et al. 1974; Germer et al. 2015).

Scleractinian taxonomically restricted genes (TRGs) also exhibited a wider variety of distribution patterns, ranging from being present across both robust and complex corals down to small set of scleractinian families only (*e*.*g*. galaxin-2 and SAPs). The former are of particular interest for the evolution of corals. Although different time estimates have been put forward, the accepted consensus places the divergence of Complexa and Robusta in the Palaeozoic, prior to the (ca. 240 Ma) appearance of fossil modern scleractinians in the early/mid-Triassic (Romano and Palumbi 1996; Stolarski et al. 2011; Chuang et al. 2017). The discovery of palaeozoic scleractinian-like fossils does support a Palaeozoic origin for the group, with consequent fossil gaps likely being caused by poor preservation or abiotic conditions hindering the deposition of skeletons (Stolarski et al. 2011). Whether a particular skeletogenic protein was available to the common ancestor of complex and robust scleractinian corals is thus of particular evolutionary interest. It allows to determine which components of the *biomineralization toolkit* preceded the Triassic appearance of the skeleton and whether putative palaeozoic scleractinians had access to the same molecular machinery currently employed by modern representatives of this group. In this regard, one biomineralization-related event that might have preceded the Complexa-Robusta divergence appears to be the expansion in the number of acidic residues within acidic proteins. The close phylogenetic relationship between P27 and B3EWY7 - which are best blast reciprocal hits - is supported by the high similarity in the location and structure of their acidic regions.

A similar scenario could also apply to galaxin *sensu stricto*. These proteins have been proposed to be independently recruited by and within scleractinians families (*e*.*g*. Pocilloporidae) (Bhattacharya et al. 2016), implying that the protein acquired its calcification-related role after the Complexa-Robusta split. However, the presence of representatives of both robust and complex corals within the galaxin *sensu stricto* clade here described does point to an alternative scenario in which galaxin recruitment for biomineralization occurred only once and prior to the divergence of these clades. The relationship between *A*. *millepora* galaxin 1 and galaxin 2 remains on the other hand uncertain due to the current lack of support in phylogenetic analyses. Despite this, phylogenetic analysis allows to confidently argue that the protein is present in the family Agariciidae and Acroporidae and it should be considered a true (*sensu stricto*) galaxin. One aspect that remains unresolved concerns the evolutionary history of galaxin *sensu stricto* outside Scleractinia. Extensive divergence between scleractinians and other cnidarians could have eroded the evolutionary signal in these proteins (Forêt et al. 2010). Nevertheless, inability to obtain supported phylogenies for galaxin proteins might also be currently exacerbated by the inclusion of several, possibly functionally diverse, galaxin-related proteins in phylogenetic analyses. Similarity between galaxin *sensu stricto* and other galaxin-related proteins is often low and restricted to di-cysteine motifs. Combined with the current lack of additional defining features for galaxins, this makes Blast-based homolog selection potentially non-optimal and can lead to the inclusion of unrelated proteins within protein datasets. Although our analysis is not immune to these limitations, expanding homolog selection beyond best-matches only in phylogenetic analysis helped to identify putative erroneous inclusions. An example described here is the *F*. *scutaria* protein 6662. When a galaxin *sensu stricto* sequence is used as a query, this sequence is the only hit in *F*. *scutaria*. Including multiple galaxin Blast matches per species did reveal however that the protein is instead a scleractinian galaxin-related protein. The presence of ‘undetected’ galaxin-related proteins, erroneously considered genuine galaxin *sensu stricto* homologs, could thus explain the previously described galaxin polyphyly (Bhattacharya et al. 2016).

Finally, in contrast to scleractinians, octocoral TRGs were found conserved across the groups showing similar distributions. Intra-Octocorallia analysis are of potential interests, as they might allow to identify differences between calcite and aragonite-deposing species, and similarities between aragonitic animals within Anthozoa (*i*.*e*. *H*. *coerulea* and Scleractinia). The presence of TRGs (such as scleritin) in species belonging to all the three major octocoral clades (McFadden et al. 2006), indicates that TRGs, although restricted to octocorals, were present in the common ancestor of the subclass. This does on one hand point towards a certain degree of commonality in spite of the different biomineralization strategies (calcite *vs*. aragonite). On the other it could be related to scenarios in which, as galaxin *sensu stricto* (Forêt et al. 2010), the protein played a different ancestral function with subsequent lineage-specific recruitments in biomineralization.

Here, we conducted a distribution and phylogenetic analysis of coral biomineralization genes to provide a comprehensive homolog mapping and fine-scaled phylogenies of selected genes. Through a broad taxon sampling, our work allowed us to detect similarities and differences between different taxonomic groups and investigate patterns of presence/absence associated with skeleton polymorph. This led to an alternative hypothesis for the evolution of galaxins and provided a detailed phylogeny of coral acidic proteins that revealed the increase of acidic residues during cnidarian evolution. We also provide information on the evolution of proteins likely involved in biomineralization, like sponge collagen IV. With the inclusion of four new octocoral transcriptomes we have closed the existing bias towards certain cnidarian taxa, specifically scleractinian corals, however gaps still exists. For instance, interesting groups like calcifying hydrozoans remain unexplored and their inclusion in future studies on biomineralization will certainly contribute to our understanding of this process in Cnidaria. Proteomic investigations of the SOM of calcifying cnidarians other than scleractinian corals and of sponges might reveal the presence of shared *skeletome* components adding support to the transcriptomic presence patterns described here, and will help discover lineage-specific innovations linked to calcification in these groups.

## Acknowledgements

SV was supported by the German Research Foundation (DFG) grant Va1146-2/1“MINORCA”. GW was supported by LMU Munich’s Institutional Strategy LMUexcellent within the framework of the German Excellence Initiative, and the German Research Foundation (DFG) grant Wo896/18-1 “MINORCA”, and from the European Union’s Horizon 2020 research and innovation programme under the Marie Skłodowska-Curie grant agreement No 764840 (ITN IGNITE). We thank the Genomics Core Facility at EMBL Heidelberg for library preparation and sequencing and Dr. Meino Rohlf, Gene Center - Dr. von Hauner Children’s Hospital, for providing access to their sequencing facilities. We thank Dr. Peter Naumann for technical assistance and maintenance of the aquaria facilities and coral culturing, as well as Simone Schätzle and Gabriele Büttner for assistance in the laboratory. SV is indebted to N. Villalobos Trigueros, M. Vargas Villalobos, S. Vargas Villalobos and S. Vargas Villalobos for their constant support.

