## Supplemental Fig. 1 for "New non-bilaterian transcriptomes provide novel insights into the evolution of coral *skeletomes*"

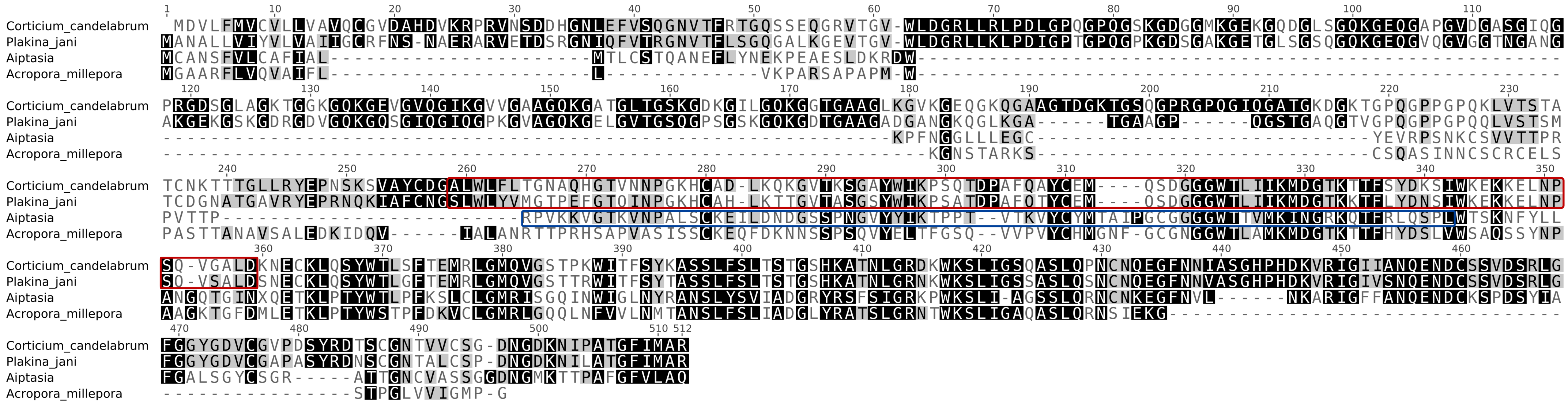


**S.Fig 1** Alignment of USOMP5-like sequences showing the position of the homologous superfamily Fibrinogen, alpha/beta/gamma chain, C-terminal globular, subdomain 1 (IPR014716) in homoscleromoprh sponges [red box] and Aiptasia [blue box]. No domain was detected in *A. millepora.*
