## Supplemental Fig. 2 for "New non-bilaterian transcriptomes provide novel insights into the evolution of coral *skeletomes*"

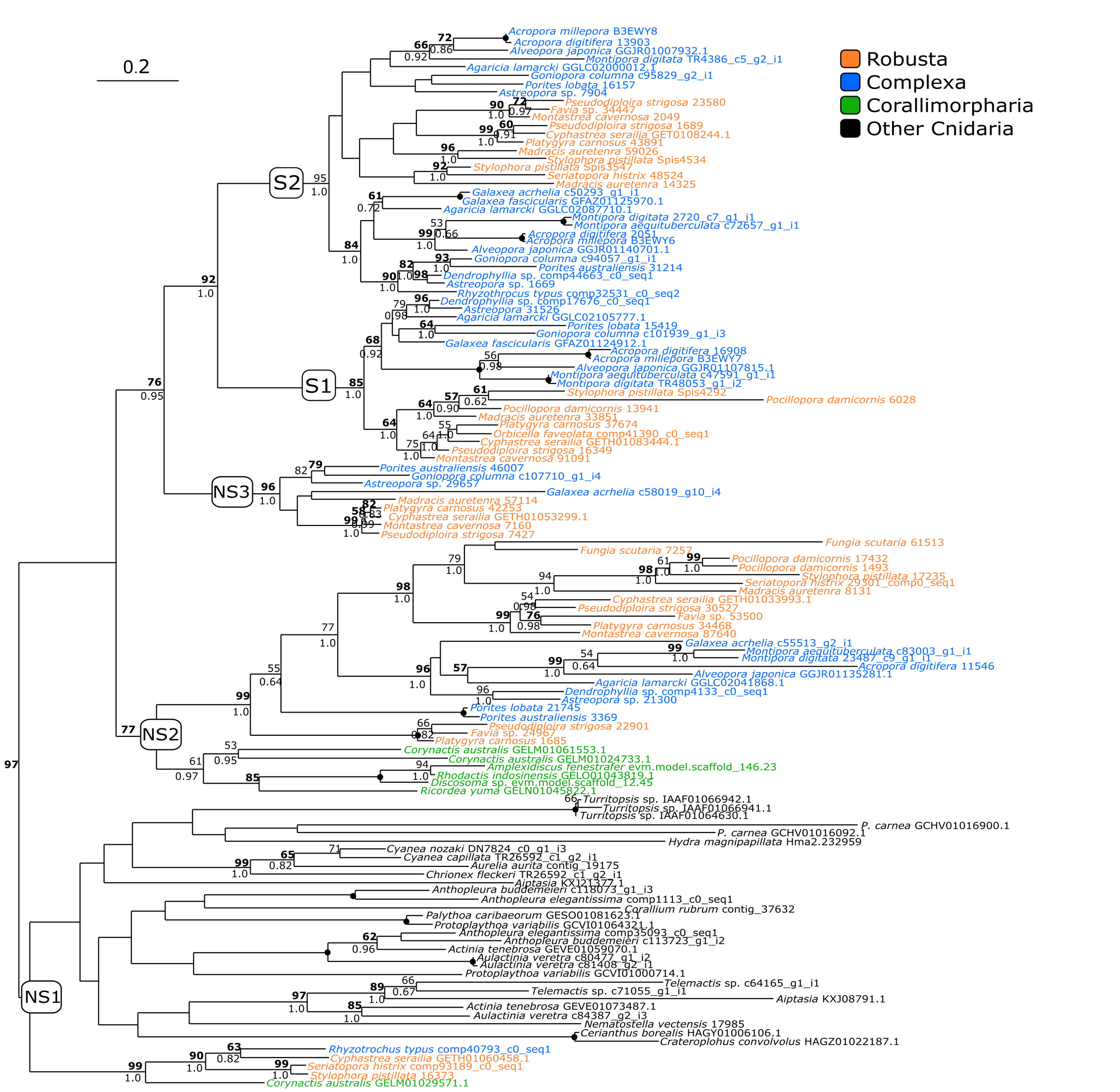


**S.Fig 2** Maximum Likelihood tree (500 bootstrap replicatesof cnidarian acidic proteins. Protein sequences aligned with MUSCLE. Best-fit model: WAG + Γ + I. Black dot on node indicates full support (100 bootstrap - 1.0 Posterior Probability. Boostrap values in bold: support is > 50 also in phylogeny based on MAFFT alignment.
