## Supplemental Fig. 3 for "New non-bilaterian transcriptomes provide novel insights into the evolution of coral *skeletomes*"

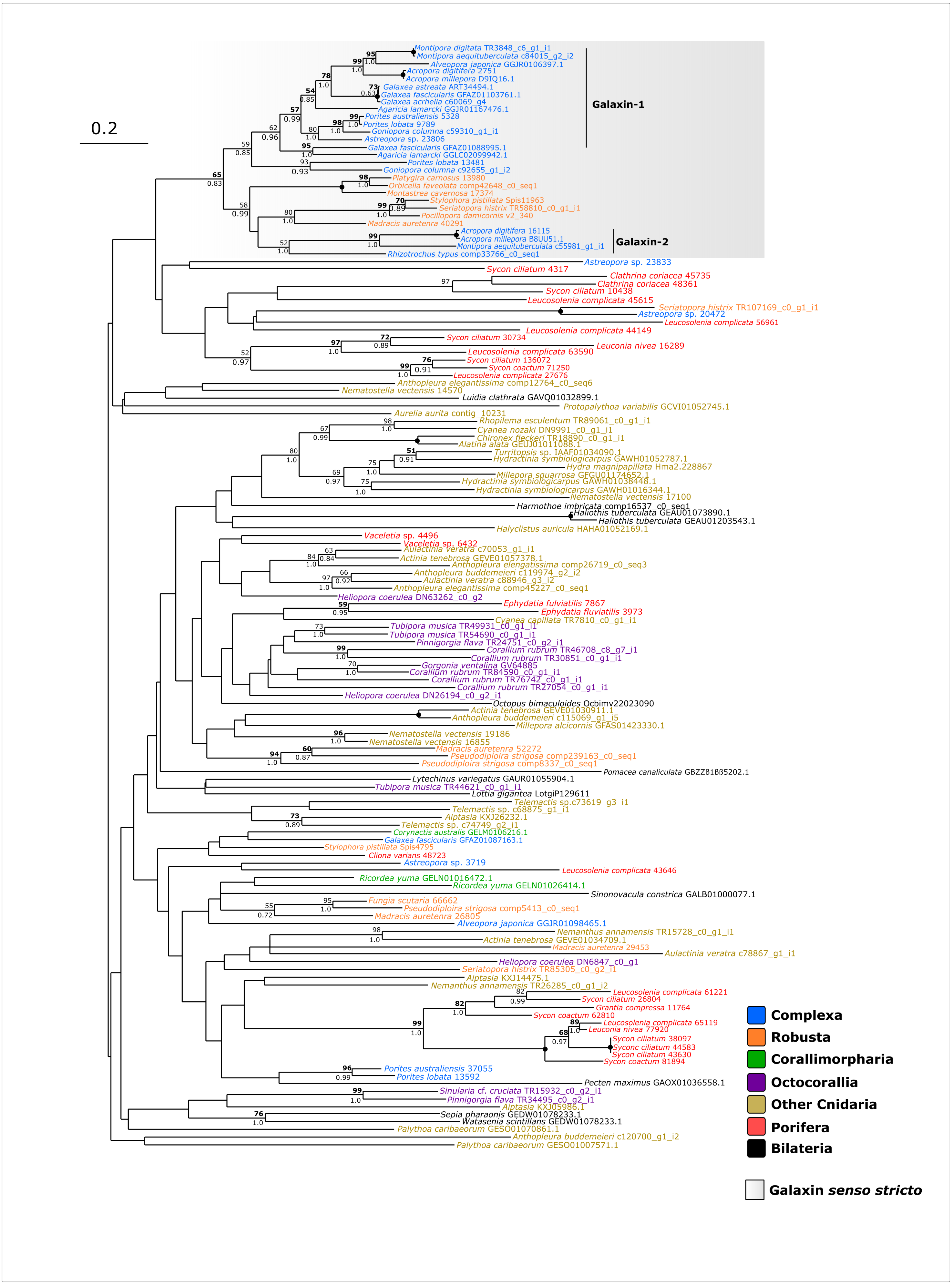


**S.Fig4** Phylogenetic analysis (500 bootstraps) of metazoan galaxin-related proteins. Tree displayed in figure based on protein sequences aligned with MUSCLE alignment. Bold number: node supported (>50) also in MUSCLE phylogeny. Dot on node indicates full support (100 bootstrap, 1.0 posterior probability) in both phylogenies. Support for nodes with bootstrap < 50 not shown regardless of posterior probability value.
