## Supplemental Fig. 4 for "New non-bilaterian transcriptomes provide novel insights into the evolution of coral *skeletomes*"

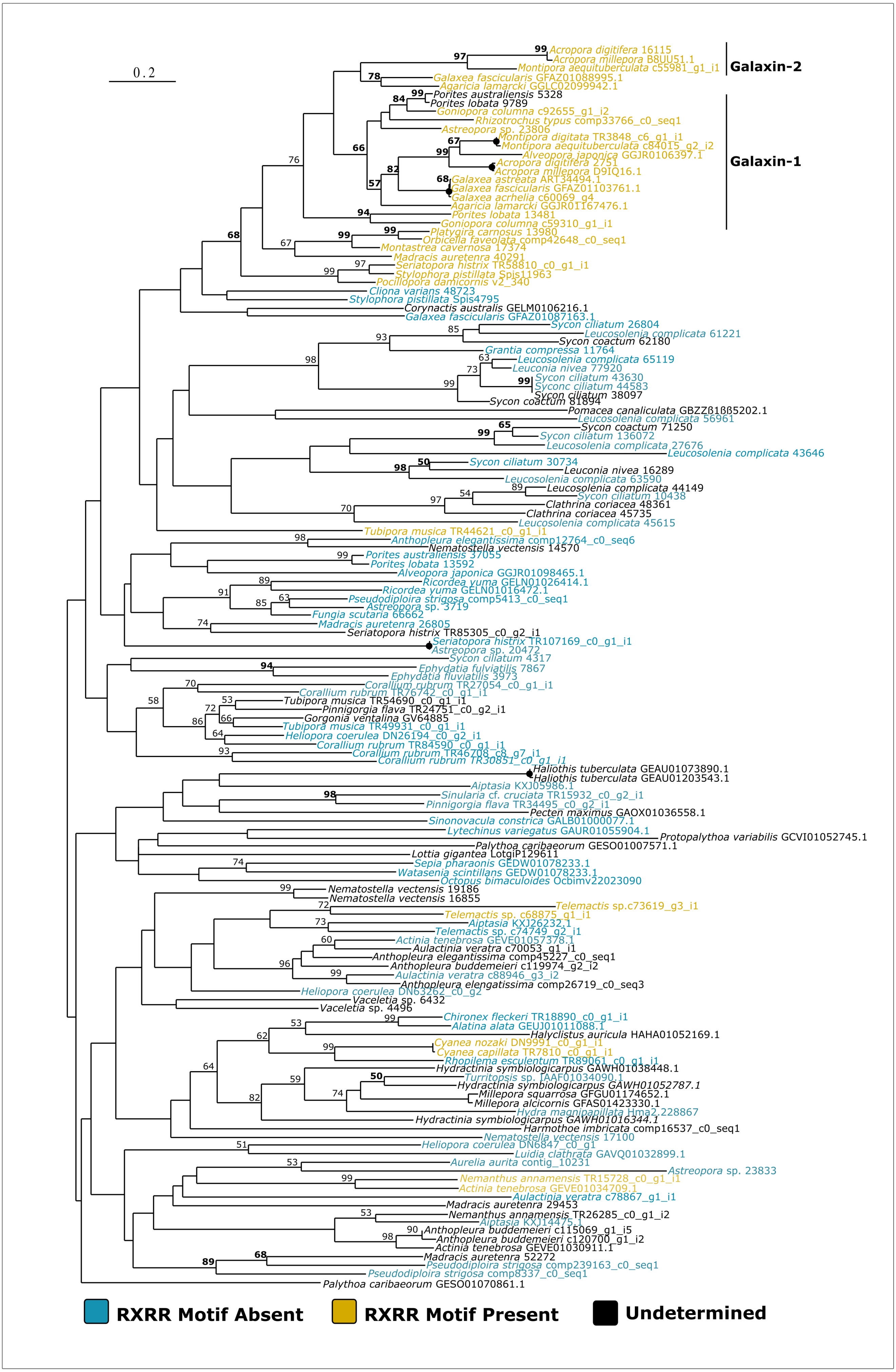


**S.Fig5** MAFFT-based phylogenetic analysis of galaxin-related proteins (Fig.3)highlighting the presence of the RXRR motif described in Fukuda et. al (2003).
