## Supplemental Fig. 8 for "New non-bilaterian transcriptomes provide novel insights into the evolution of coral *skeletomes*"

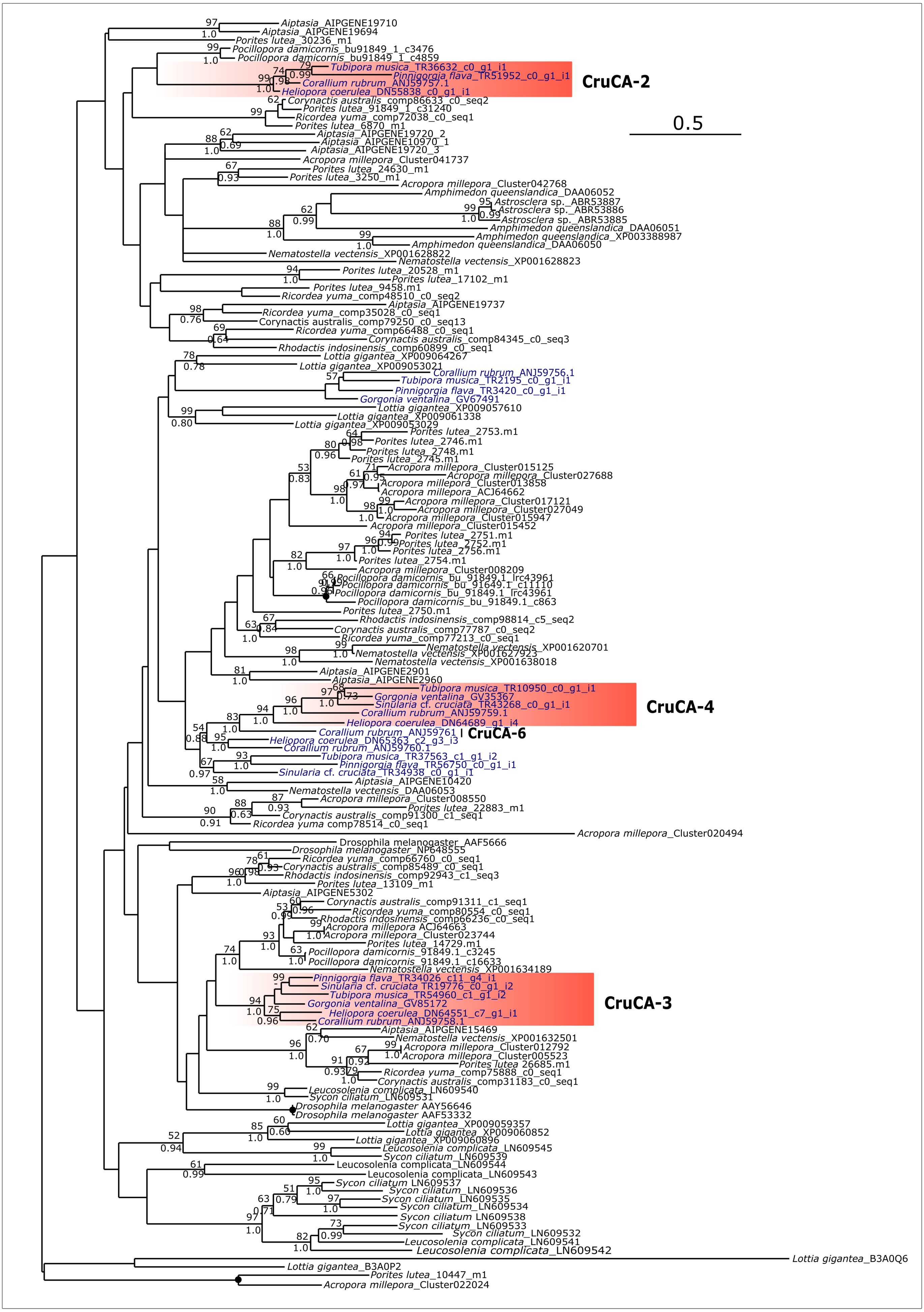


**S.Fig3** Phylogenetic analysis (500 bootstraps) of octocoral carbonic anhydrases (CAs) (MUSCLE). Octocoral CAs are in blue.Sequences were added to the dataset used inLin et al. (2017). Best-fit model: LG + G. Black dot on node indicatesfull support (100 relative bootstrap - 1.0 posterior probability).
