## Supplemental Fig. 5 for "New non-bilaterian transcriptomes provide novel insights into the evolution of coral *skeletomes*"

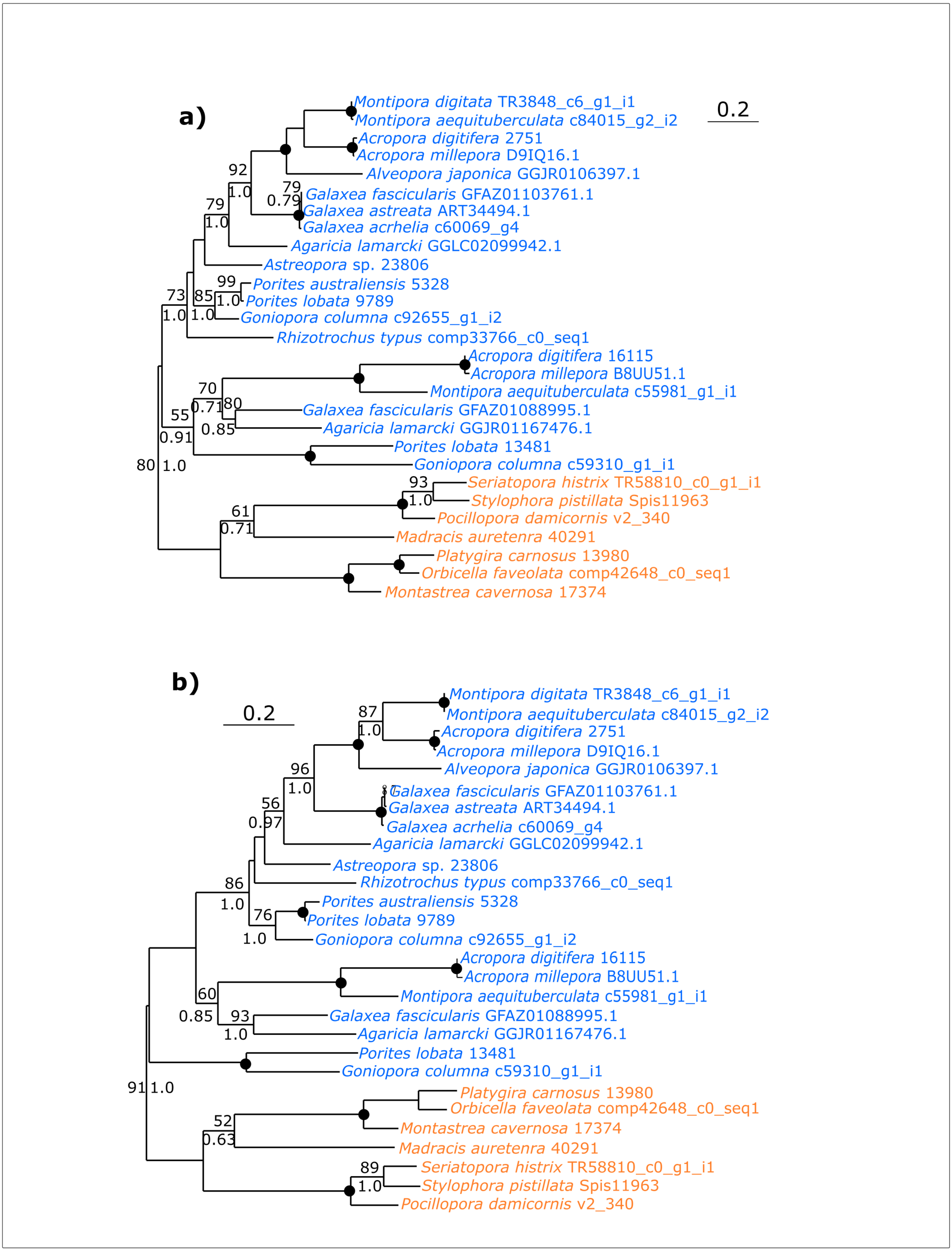


**S.Fig 3** Phylogenetic analysis (500 bootstrap replicates) of galaxin *senso stricto* based on MUSCLE (**a**) and MAFFT (**b**) alignes sequences. Best-fit model for both alignments: JTT + Γ + I. Black dot on node indicates full support (100 bootstrap and 1.0 posterior probability. Maximum-likelihood and bayesian analysis were performed with PhyML 3.1 (in Seaview 4) and MrBayes 3.2.6 respectively. For the latter a burn-in fraction of 20% was applied.
