## Supplemental Fig. 9 for "New non-bilaterian transcriptomes provide novel insights into the evolution of coral *skeletomes*"

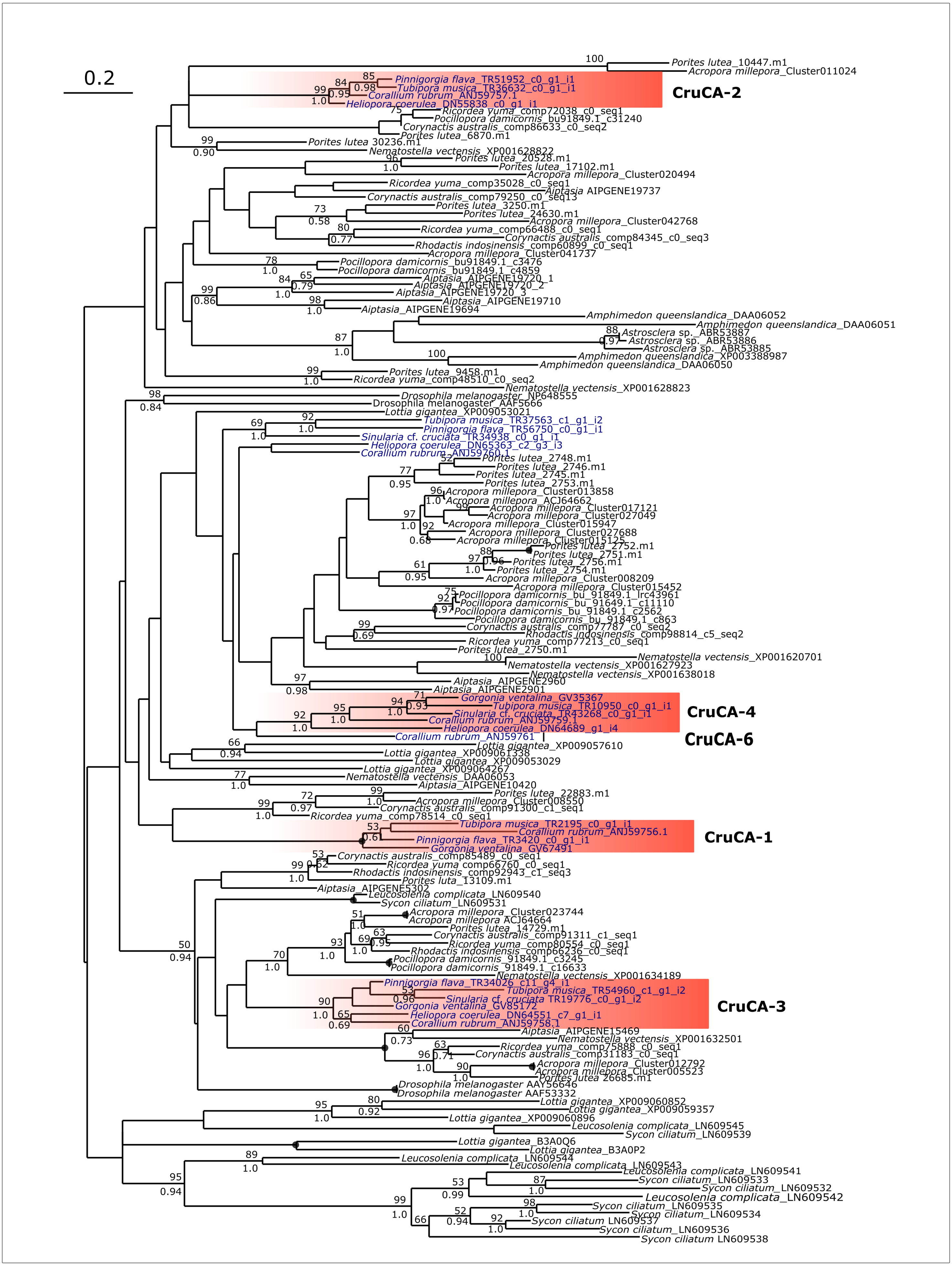


**S.Fig2** Phylogenetic analysis (500 bootstraps) of octocoral carbonic anhydrases (MAFFT). Octocoral CAs are in blueSequences were added to the dataset used in Lin et al. (2017). Best-fit model: LG + G. Black dot on node indicates full support (100 relative bootstrap - 1.0 posterior probability.
