## Supplementary figures and images for "New non-bilaterian transcriptomes provide novel insights into the evolution of coral *skeletomes*"

### Supplemental Fig. 6

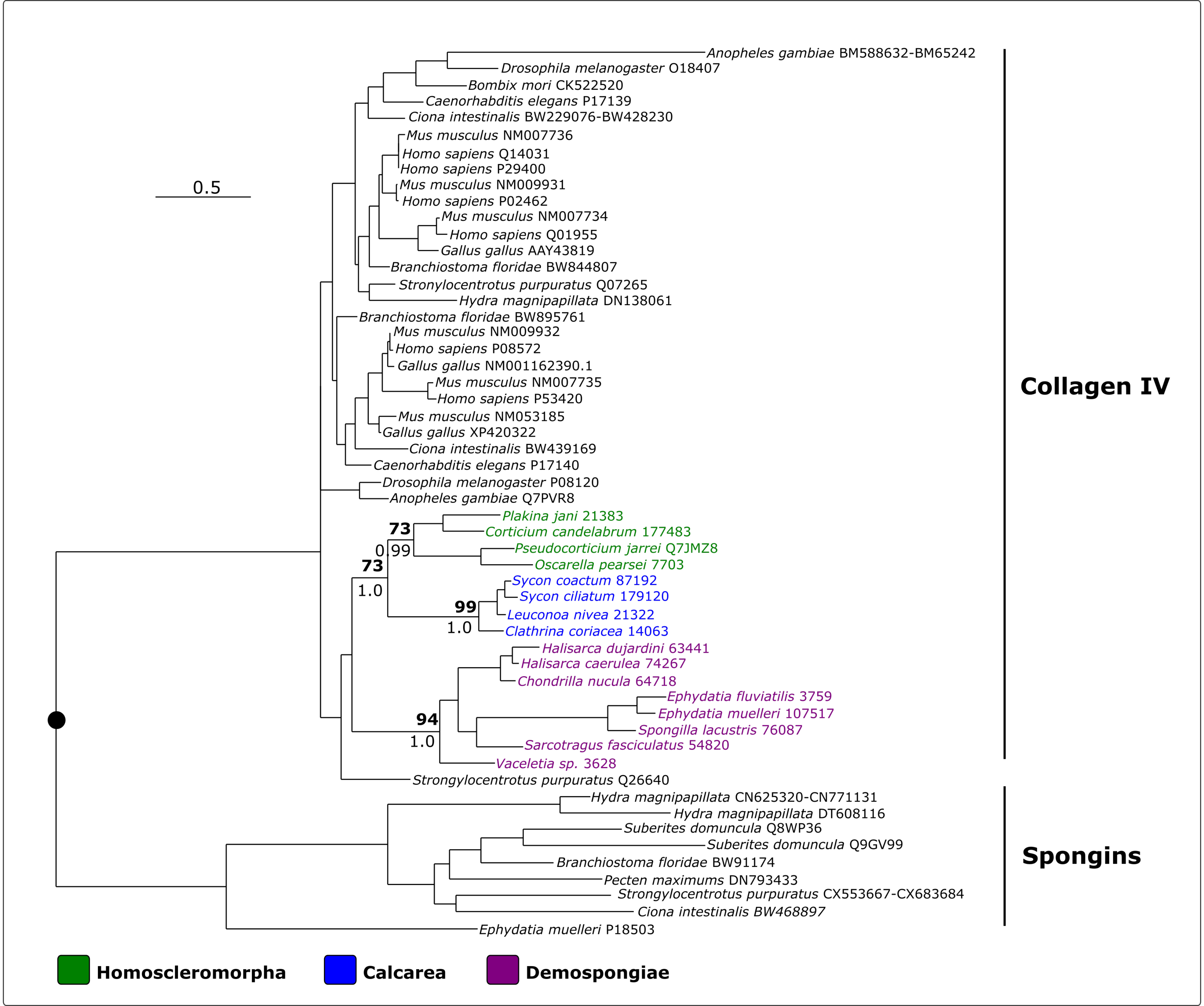

### Supplemental Fig. 7

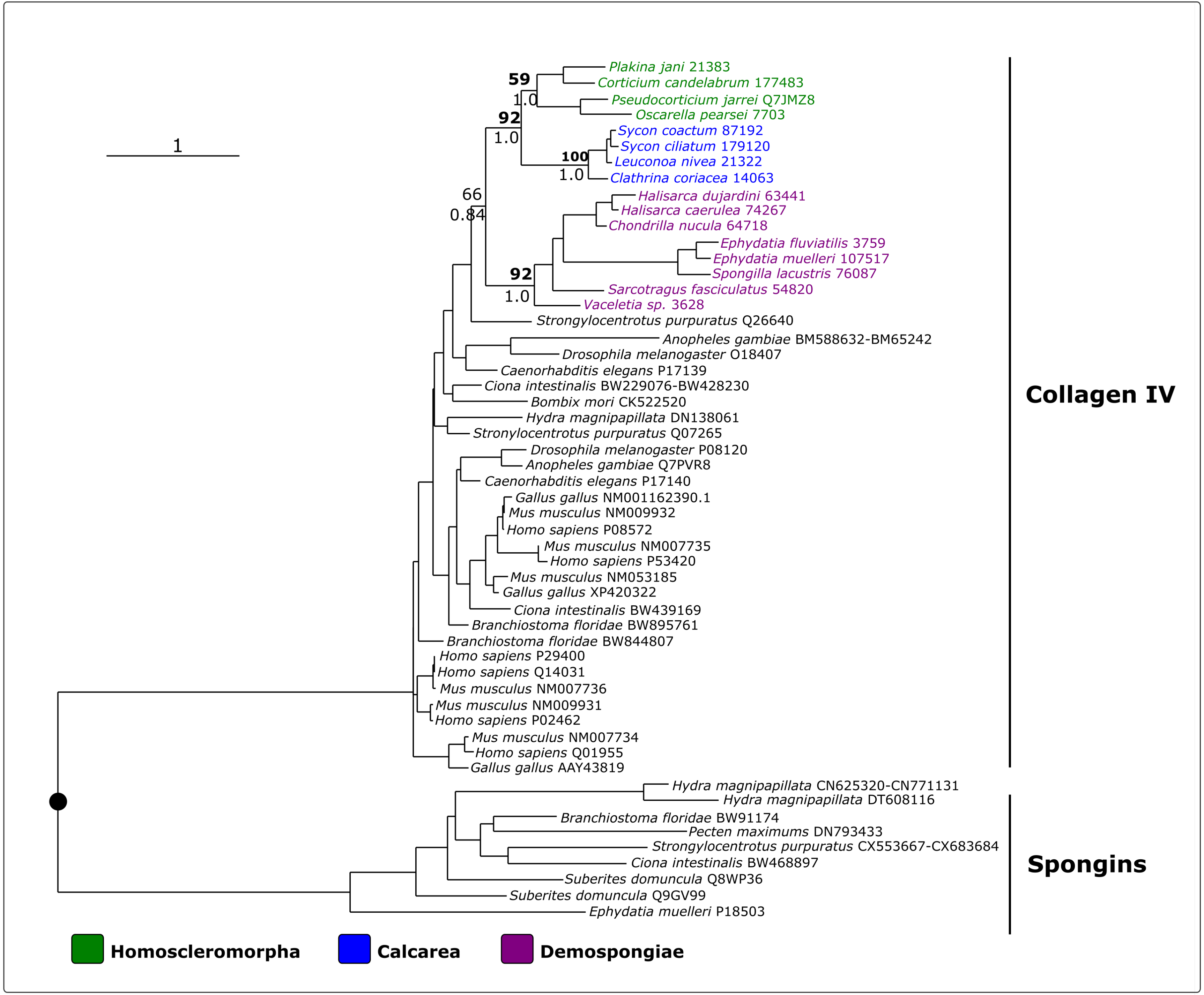
